# Viral expansion after transfer is a primary driver of influenza A virus transmission bottlenecks

**DOI:** 10.1101/2023.11.19.567585

**Authors:** Katie E. Holmes, David VanInsberghe, Lucas M. Ferreri, Baptiste Elie, Ketaki Ganti, Chung-Young Lee, Anice C. Lowen

## Abstract

For many viruses, narrow bottlenecks acting during transmission sharply reduce genetic diversity in a recipient host relative to the donor. Since genetic diversity represents adaptive potential, such losses of diversity are thought to limit the opportunity for viral populations to undergo antigenic change and other adaptive processes. Thus, a detailed picture of evolutionary dynamics during transmission is critical to understanding the forces driving viral evolution at an epidemiologic scale. To advance this understanding, we used a barcoded virus library and a guinea pig model of transmission to decipher where in the transmission process influenza A virus populations lose diversity. In inoculated guinea pigs, we show that a high level of viral barcode diversity is maintained. Within-host continuity in the barcodes detected across time furthermore indicates that stochastic effects are not pronounced within the inoculated hosts. Importantly, in both aerosol-exposed and direct contact animals, we observed many barcodes at the earliest time point(s) positive for infectious virus, indicating robust transfer of diversity through the environment. This high viral diversity is short-lived, however, with a sharp decline seen 1-2 days after initiation of infection. Although major losses of diversity at transmission are well described for influenza A virus, our data indicate that events that occur following viral transfer and during the earliest stages of natural infection have a central role in this process. This finding suggests that host factors, such as immune effectors, may have greater opportunity to impose selection during influenza A virus transmission than previously recognized.

## INTRODUCTION

The high mutation rate characteristic of RNA viruses [1, 2], coupled with genetic recombination [3, 4] and/or reassortment [5] of segmented genomes, enables constant production of viral variants [6–8]. In turn, this variation provides the substrate for viral evolution [9–11]. For some viruses, including influenza A virus and SARS-CoV-2, viral evolution and viral spread through host populations occur on a similar timescale, such that each shapes the other [12]. Understanding viral evolution is therefore crucial in our efforts to control outbreaks.

Selection and genetic drift are two major drivers of evolutionary change in viral populations [13]. Selection is a deterministic process whereby differences in fitness result in changes in variant frequencies over time. In contrast, genetic drift is a stochastic process whereby changes in variant frequencies result from chance. In general, when population sizes are large, selection predominates over drift and changes in variant frequencies result from differences in the variants’ reproductive output. When population sizes are small, drift instead predominates and changes in variant frequencies result simply from random chance associated with reproductive output. Both selection and drift can reduce genetic diversity, but by different means. Under drift, the loss of a variant comes about as a chance event. Under selection, the loss of a variant comes about from the variant having lower fitness. As such, these processes have different impacts on a population’s fitness trajectory: if driven by selection, the loss of diversity increases viral population-level fitness and as such results in viral adaptation. If driven by drift, the loss of viral diversity could (by chance) either increase or decrease population-level fitness. For influenza viruses, the dynamic interplay between selection and genetic drift is still poorly understood [13–15].

A reduction in viral population size during transmission has been documented for many viruses [16–20]. Termed a transmission bottleneck, this effect is characterized by the establishment of a new infection by few viruses derived from a large source population. The reduction in population size is typically accompanied by a loss of population diversity, known as a genetic bottleneck. Classically, bottlenecks are defined as random events and are therefore a source of genetic drift [21]. When a random bottleneck is active, even highly adaptive variants may go extinct. In the case of HIV, however, the sharp reduction in population size at transmission is a selective process favoring variants that use CCR5 co-receptors [22–24]. The term selective bottleneck is often used to describe this deterministic process.

As for other viruses, loss of diversity during influenza A virus transmission has been documented. In humans with naturally occurring infections, as few as one to two viral genomes from an infected individual established in an uninfected individual [13]. Studies using influenza A virus populations with inserted genetic barcodes similarly revealed narrow transmission bottlenecks in animal models [25, 26]. More relaxed bottlenecks have been reported in natural hosts aside from humans, suggesting roles for host biology and/or modes of transmission in shaping species-specific between-host dynamics [27, 28]. Finally, avian influenza A virus transmission in mammals has been shown to involve a tight bottleneck with reductions in viral diversity attributed to either stochastic or selective forces [29–31].

While it is known that diversity is lost during influenza A virus transmission, it is unclear at which point – in the donor host, the environment, or the recipient host – this takes place. The specific stage at which diversity is lost is likely to define the potential for selection to act during transmission. We therefore made an influenza A virus library with 4096 potential barcodes to monitor the fate of many unique viral lineages within infected guinea pigs and through the course of transmission to contacts. Our data reveal that viral diversity remains high in inoculated animals throughout infection, and many viral genomes are transferred to animals exposed by aerosols or direct contact. However, a severe bottleneck occurs in exposed animals 1-2 days after the initiation of infection, such that few lineages are sustained in the context of population expansion. Thus, in our system, losses of diversity are primarily driven by events that occur during the expansion of infection, not during the process of donor-to-recipient transfer itself. Importantly, this phase of expansion could be a previously unrecognized opportunity for selection to operate.

## RESULTS

### Generation of a Barcoded Influenza A Virus Library

We designed a genetic barcode for influenza A/Panama/2007/99 (H3N2) virus (Pan/99) with the goals of producing a diverse viral population while avoiding attenuation of viral replication and fitness differences among barcode variants. To achieve high diversity, twelve nucleotide sites within a 50-nucleotide region of the NA segment were made bi-allelic. Since one of two bases was possible at each of 12 positions, 2^12^ = 4096 unique barcodes were introduced (**Figure 1A**). To avoid attenuation, we introduced polymorphisms within the native sequence of Pan/99 rather than inserting a foreign sequence. To avoid fitness differences among the barcode variants, the polymorphisms introduced were synonymous and chosen based on their natural occurrence in H3N2-subtype influenza A viruses circulating in humans from 1994 to 2004. We reasoned that naturally occurring variants detected in circulation would be likely to have minimal fitness effects. We named the gene segment modified in this way Pan/99 NA-BC.

**Figure 1.**
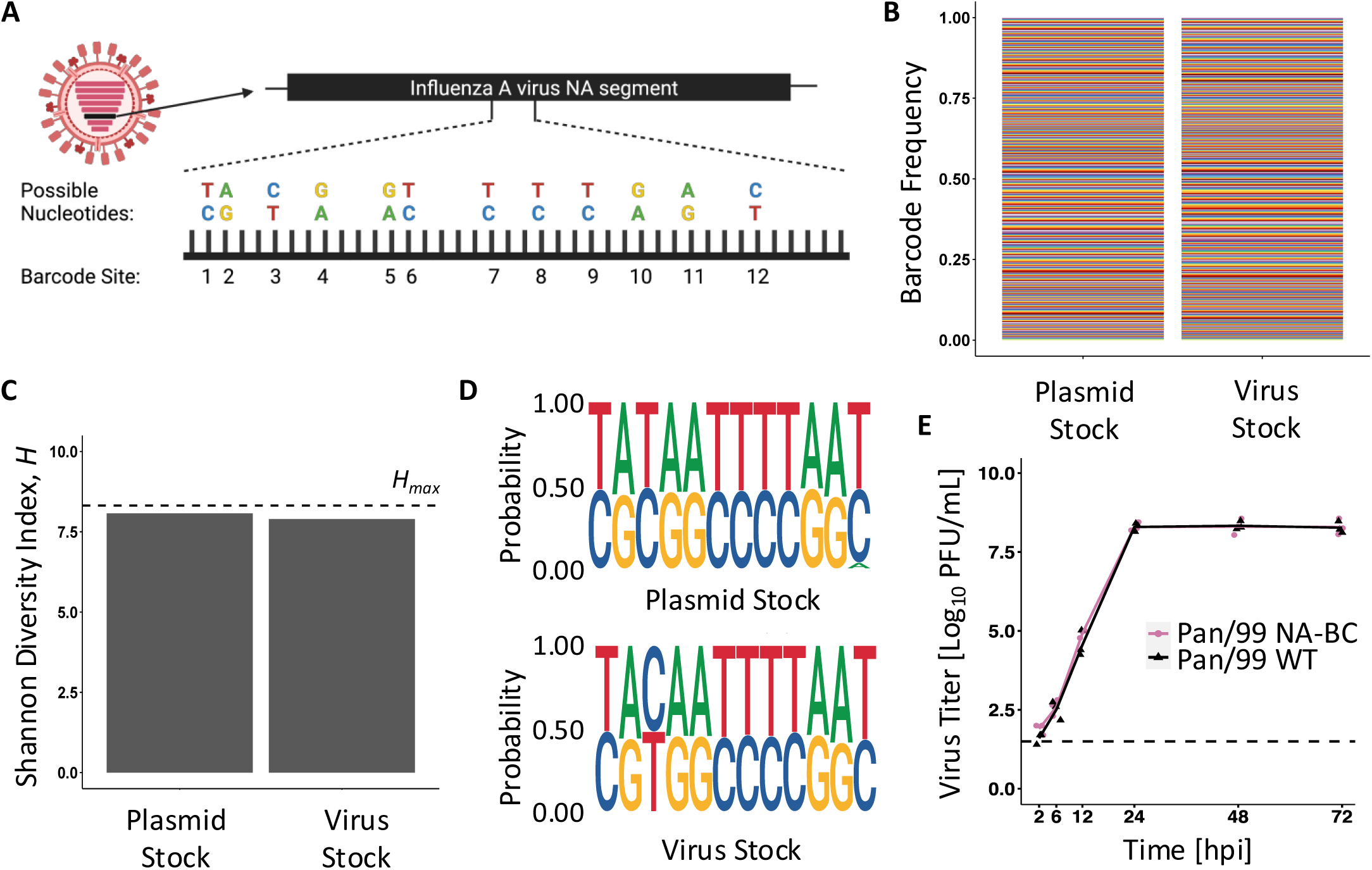
NA barcode diversity is maintained in both plasmid and virus stocks, and the barcode does not affect overall fitness. **A)** Barcode design for the NA segment of influenza A/Panama/2007/99 (H3N2) virus. At each of twelve sites, one of two nucleotides is possible, allowing for up to 4096 unique barcodes within the population. **B)** Barcodes detected in the pDP Pan/99 NA-BC plasmid preparation and passage 1 stock of Pan/99 NA-BC virus. Colors represent unique barcodes, and their frequencies within the stock are indicated by the height of the color. **C)** Shannon Diversity Index (*H*) of the stock samples is compared to the maximum possible diversity for this system (*H_max_* = 8.32, shown with a horizontal dashed line). This theoretical maximum reflects a population in which all 4096 potential barcodes are equally represented. **D)** Sequence logo plots demonstrate the sequence motifs present in plasmid and passage 1 virus stocks. Each pair of nucleotides represents one of the twelve bi-allelic sites. The height of the letter indicates the corresponding nucleotide frequency in the stock. **E)** Multicycle replication of Pan/99 WT (black) and Pan/99 NA-BC (pink) viruses. Infections were performed in triplicate, and data points show results from individual cell culture dishes. Lines connect mean titers. Dashed line indicates limit of detection (50 PFU/mL). Two-way ANOVA showed no significant difference between viruses at any time point (*p* = 0.11). Figure 1A created in BioRender.

To evaluate barcode diversity in the reverse genetics plasmid and the passage 1 (P1) viral stock carrying Pan/99 NA-BC, we subjected the barcode region to next-generation sequencing. No barcode was found to be dominant in the plasmid library or the virus stock, and no nucleotide was dominant at any site within the barcode (**Figure 1B, D**). An aberrant nucleotide was detected at the twelfth barcode site in the plasmid preparation but was not carried through to the virus stock. To quantify diversity, we applied the Shannon Diversity Index (*H*), which considers the richness (i.e., the number of species present) and the evenness (i.e., the abundance of the present species) in a community [32]. To calculate *H*, each unique barcode was taken to represent a species. The measured diversity in the plasmid and virus stocks was 8.07 and 7.90, respectively (**Figure 1C**), near to the theoretical maximum (*H_max_*) for this system of 8.32 and revealing little loss of barcode diversity during viral recovery from cDNA. To determine whether the barcode in the NA segment altered fitness of the Pan/99 virus, multi-cycle replication was evaluated in MDCK cells. No significant differences between wild type and Pan/99 NA-BC viruses were detected (**Figure 1E**).

### Modeling Transmission of Pan/99 NA-BC Virus in Guinea Pigs

To evaluate viral dynamics within and between hosts, we inoculated guinea pigs with 5 x 10^4^ plaque-forming units (PFU) of Pan/99 NA-BC virus intranasally [33]. After 24 hours, we placed a naïve animal with each inoculated animal in either direct or aerosol contact (**Supplemental Figure 1A**). Three independent experiments were performed, each including four direct contact and four aerosol transmission pairs. Viral shedding, assessed by plaque assay of daily nasal lavage samples, was similar for all inoculated animals (**Supplemental Figure 1B**). Transmission efficiency varied from 75% to 100% for pairs in direct contact and from 50% to 100% for pairs in aerosol contact. The daily nasal lavages collected from both inoculated and exposed animals furnished samples with which to investigate viral population dynamics and the drivers of the influenza A virus transmission bottleneck.

### Stochastic Effects are Not Pronounced in Inoculated Animals

To determine how barcode diversity changed during infection in inoculated animals, we evaluated the barcodes present in nasal lavage samples by next-generation sequencing. In all directly inoculated animals, many barcodes were detected throughout the course of infection (**Figure 2** and **Supplemental Figure 2**). Using the Shannon Diversity index (*H*), we found that barcode diversity either remains consistent or declines slightly over time in inoculated animals (**Figure 3** and **Supplemental Figure 3**). Changes in diversity are furthermore mirrored by changes in both barcode richness and evenness, indicating that declines in diversity are driven both by loss of barcode species from the within-host population and skewing of the relative frequencies of individual barcodes that comprise the population (**Figure 3** and **Supplemental Figure 3**).

**Figure 2.**
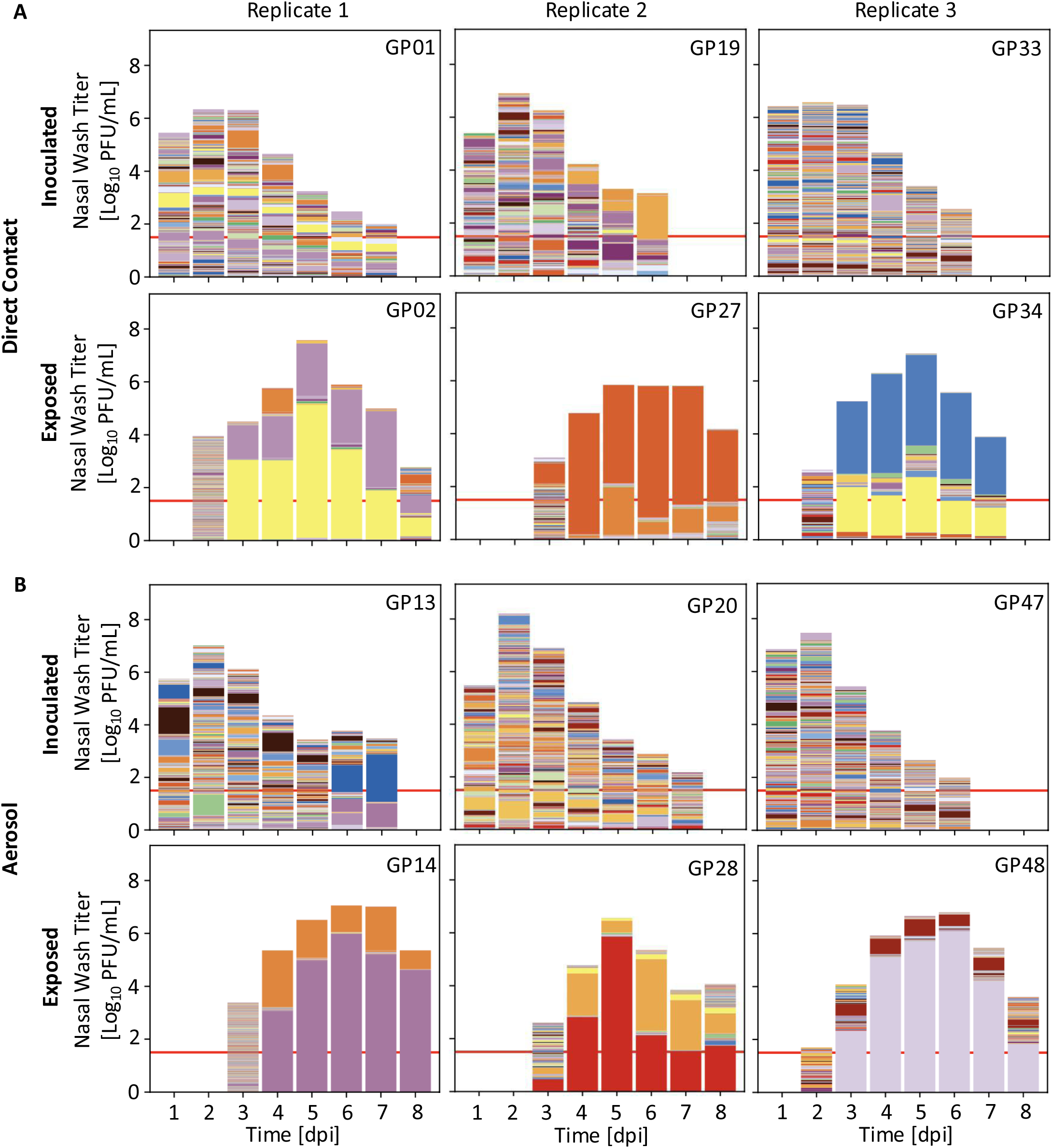
Population diversity declines between inoculated and exposed guinea pigs. Viral titers in nasal lavage samples are indicated by the overall height of the bar. Red lines show LOD (50 PFU/mL). Colors within the bars represent unique barcodes, and the height of each color indicates barcode frequency within the sample. Plots for individual animals are paired with those of their cage mate. Representative pairs for direct contact (**A**) and aerosol (**B**) exposure are shown from three experimental replicates. Guinea pig ID numbers are shown in the upper right corner of each plot. Data from additional animals are shown in Supplemental Figure 2.

**Figure 3.**
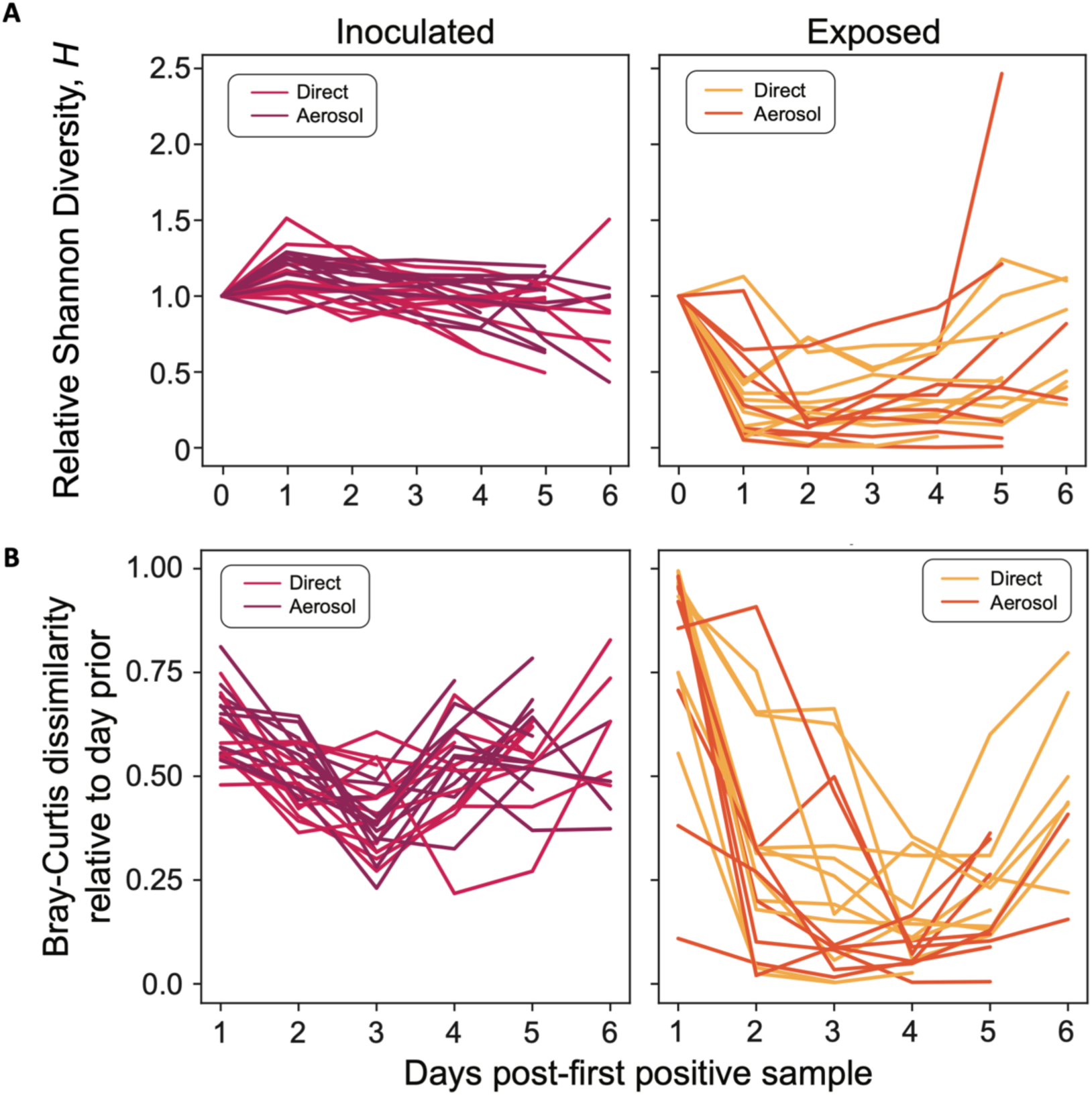
Initial high barcode diversity in exposed animals plummets after the first 1-2 days of viral positivity. **A)** For each exposed animal, the Shannon diversity index *(H*) was normalized to that on the first day of viral positivity (as determined by plaque assay). Data from a total of 24 inoculated animals (left) and 19 exposed animals (right) across three experiments are shown. The Shannon diversity in each animal relative to their first day of positivity is on average 3.7x lower in exposed animals compared to inoculated animals across all timepoints (p<0.0001). **B)** Changes in barcode composition between successive days was assessed using the Bray-Curtis dissimilarity index to compare barcode composition in each sample to that observed on the prior day. Lines connect data points for one animal, with inoculated (left) and exposed (right) animals shown on separate graphs.

### High Initial Diversity in Exposed Animals Precedes a Sharp Decline

To determine how barcode diversity changed between donor and recipient animals, barcodes present in nasal lavage samples of recipients were analyzed. Strikingly, at the earliest timepoint(s) positive for infectious virus in recipients, many barcodes are present, signifying transfer of an appreciably large and diverse viral population through the environment (**Figure 2** and **Supplemental Figure 2**). This observation is true of both aerosol-exposed and direct contact animals. Populations present early in exposed animals are characterized by high barcode diversity, richness, and evenness (**Figure 3** and **Supplemental Figure 3**).

However, the observed high barcode diversity in exposed animals is short-lived, with a sharp decline seen within 1-2 days of the initiation of infection (**Figure 2, Supplemental Figure 2, and Supplemental Figure 3**). Both richness and evenness contribute to the precipitous drop in diversity seen within exposed animals (**Figure 3A** and **Supplemental Figure 3**). The stark change in population composition is unique to early times after transmission: pairwise comparisons between successive time points show high dissimilarity of barcode composition between the first and second positive days but low dissimilarity between successive days thereafter (**Figure 3B**). While the few viral lineages that penetrate the bottleneck persist through the remainder of the acute infection, those that persist differ across animals (**Figure 2** and **Supplemental Figure 2**), excluding the possibility that some barcodes carry a fitness advantage. Notably, barcode diversity in some exposed animals rebounds late in the infection.

To assess the potential for the observed population dynamics to be driven by selection acting on *de novo* mutations outside the barcoded region, we performed whole-genome sequencing of a subset of nasal lavage samples. Specifically, viral genomes collected on the first or second day of positivity in exposed animals from experimental replicates 1 and 2 were sequenced in full. Among these samples, a single variant was found above a 1% frequency, which was a nonsynonymous change present at 4.9% in the NP segment of guinea pig 31 on day 5 post-inoculation. This variant resulted in a methionine to valine substitution at amino acid 238. Thus, reductions in viral diversity observed in exposed animals were not driven by selective sweeps of *de novo* variants.

As a means of validating results obtained from targeted sequencing of the barcoded region of the viral genome, we analyzed the whole-genome sequencing reads that spanned this region. This analysis relied on reads spanning the entirety of the barcode, which were relatively rare. Nevertheless, high-level observations could be confirmed: first-positive time points showed multiple barcodes present and subsequent time points revealed the same predominant barcodes in both sequencing datasets (**Supplemental Figure 4A**). Furthermore, results of targeted barcode sequencing were validated by sequencing a subset of samples multiple times to ensure reproducibility (**Supplemental Figure 4B, C**).

To evaluate whether barcodes detected on the first day of positivity in recipient animals were carried by viable viruses, we performed plaque assays with nasal lavage samples from the first and second positive days from a subset of eight recipient animals. The barcodes present in plaque isolates were then identified by sequencing. In three of the eight animals, plaques from the first day of positivity yielded a diverse set of barcodes, indicating that many independent viral lineages present early in these three animals were represented in the infectious viral population (**Figure 4**). In the remaining five animals, the plaque isolates from the first positive day were low diversity and largely matched those derived from the second positive day (**Figure 4**).

**Figure 4.**
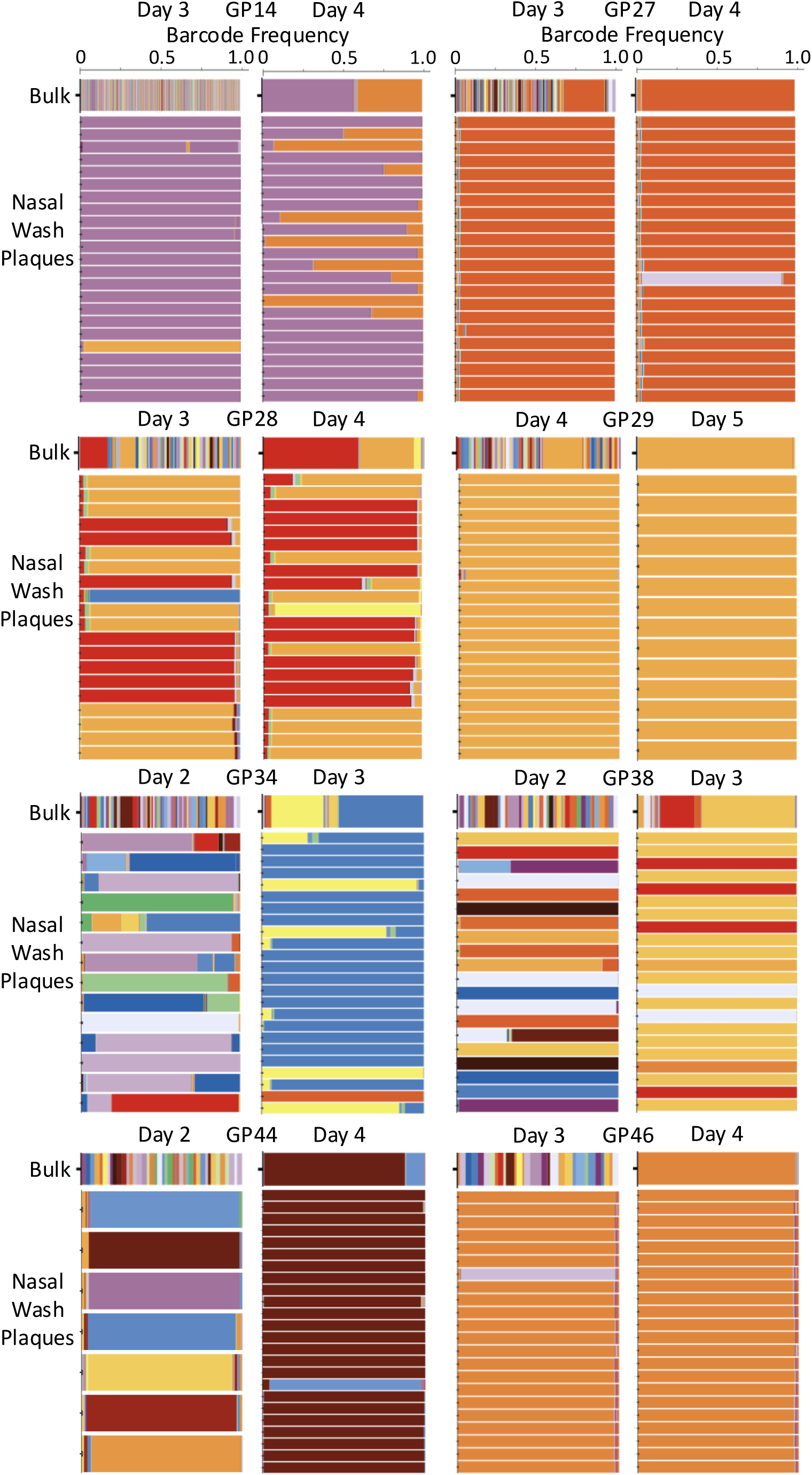
Infectious viruses isolated early after transmission carry diverse barcodes in a subset of animals. For eight exposed animals, up to 24 plaques were isolated from nasal wash samples collected on the first and second days of positivity. Barcodes detected in isolated plaques are shown beneath those detected in the corresponding nasal lavage sample, with each row representing a plaque. Each color represents a unique barcode, with frequency shown on the horizontal axis.

To test whether the differing population dynamics between donor and recipient animals relate to viral dose, we evaluated dynamics of barcode composition in animals inoculated with escalating doses of virus (**Supplemental Figure 5A**). Three animals were inoculated intranasally with 1 x 10^2^ PFU, 3.3 x 10^2^ PFU, 1 x 10^3^ PFU, 3.3 x 10^3^ PFU, 1 x 10^4^ PFU, or 5 x 10^4^ PFU of Pan/99 NA-BC virus. We saw that, at each inoculation dose, measured values for diversity, richness, and evenness are below what would be expected if all inoculated barcodes were recovered in nasal lavage samples (**Supplemental Figure 5B**). Thus, a genetic bottleneck acts within animals infected through inoculation. This bottleneck is of comparable stringency across the dose range tested. A bottleneck acting during the establishment of infection was not, however, observed in inoculated animals as it was transmission recipients.

## DISCUSSION

We used a barcoded influenza A virus system to generate a high-resolution view of changes in the composition and diversity of viral populations as they expand, contract, transmit, and establish infection in a new host. Our data suggest that viral evolutionary dynamics through a transmission event comprise two distinct stages: inter-host transfer and intra-host establishment. The bottleneck acting during the first stage can be loose, allowing many genotypes to pass to the new host. Conversely, the bottleneck acting at the second stage is tight, such that few genotypes dominate the established within-host population. The evolutionary implications of this two-stage process are potentially great. The existence of a loose bottleneck between hosts could enable selection to act efficiently on the transferred viral population *prior to* a stochastic contraction of diversity imposed by the process of viral establishment. Indeed our data offer a potential solution to a long-standing conundrum in evolutionary virology, that of disparate evolutionary dynamics at the individual and global population scales [34].

The dynamics we observe here for influenza A virus transmission are analogous to those seen in mice inoculated orally with a barcoded coxsackievirus, wherein high initial barcode diversity was followed by a sharp decline during enteric infection [35]. The observed reduction in diversity between influenza virus populations in donor animals and those that become established in recipients is furthermore consistent with prior studies on influenza in human cohorts and experimental animals [13, 25, 26, 29]. Our conclusion that early viral dynamics in the recipient make a major contribution to the loss of diversity during transmission is, however, novel. This pattern was likely not apparent in prior work due to the time points analyzed. In human cohort studies, sample collection is typically triggered by the onset of symptoms, such that the very early stages of infection are often not sampled [13]. Similarly, previous studies of the influenza A virus transmission bottleneck in experimental models did not characterize viral populations present early after transmission [25, 29].

The very transient nature of early, diverse, viral populations is also apparent within our own data sets. Direct sequencing of viral genomes sampled from the nasal cavity of recipient hosts revealed a collapse of barcode diversity over a period of approximately 24 hours. Conversely, sequencing of infectious isolates derived from the sampled populations suggested a more rapid extinction of viral lineages. Among eight recipient animals that each showed a high diversity of genomes in early nasal lavages, only three showed a high diversity of infectious viruses in these sampled populations. In the remaining five, the infectious viral populations detected early matched the low diversity populations that persisted through the course of infection. We postulate that the discrepancy seen in these five samples relates to the detection of viral genomes within infected cells when sequencing bulk nasal lavage. In line with the heterogeneity of viral burst size at a single-cell level [36–38], many infected cells will not produce progeny viruses that can be detected by plaque assay. Thus, these five samples appear to have captured a time point in which many unique barcodes were being replicated within infected cells, but the bottleneck had already occurred at the level of the released viral population. Conversely, the three samples that showed concordance between bulk nasal lavage and plaque isolates captured an earlier phase of the transition from high to low diversity.

Based on our results, we propose a model in which many viruses are delivered to the recipient and initiate infection in the nasal cavity, such that their genomes are replicated to a level detectable by sequencing. Among this first generation of infected cells, only a subset produces large bursts of progeny viruses early [36–39]. The genotypes replicated in those cells initiate a second round of infection, which is subject to similar heterogeneity. Within these very first generations of viral amplification, the few genotypes replicated rapidly and to high levels quickly dominate both infected-cell and released-virus populations. The eventual death of producer cells precipitates loss of barcode diversity in the infected-cell population. Among released viruses, competition for new target cells leads to the extinction of many minor lineages [40, 41]. Innate antiviral immunity likely hastens the contraction of barcode diversity both in infected cells and in released viruses [42, 43]. Within these contractions, some minor lineages may persist owing to long-lived cells [44, 45] or spatial isolation [46–48]. Our data suggest that such persistent lineages can expand late in infection as dominant lineages are cleared, yielding a transient rebound in barcode diversity.

In contrast to recipient animals infected through transmission, we found only modest reductions in diversity in donor animals during infection. This finding extended to animals inoculated with low doses, where a bottleneck acting at inoculation was apparent, but a growth-induced bottleneck acting during the establishment of infection was not. Since the kinetics of viral expansion may differ with infection route, this difference may be attributable to the timing of sampling. Alternatively, viral population dynamics may differ between animals infected through inoculation and transmission owing to differences in the mode of viral deposition. For example, viral aggregation state may differ, which would modify the potential for coordinated infection and, in turn, alter the extent of heterogeneity in viral burst size [38, 49]. Alternatively, or in addition, the size of the epithelial area across which viruses are delivered may differ. If this area is greater following intranasal instillation, the ensuing spatial separation of initial sites of infection may allow more lineages to establish within the host [46–48].

Seasonal influenza virus evolution at large geographic and temporal scales is characterized by a clear pattern of positive selection: on a recurring basis, antigenically distinct variants sweep the global viral population, driving epidemic spread [50–52]. In contrast, the within-host evolutionary dynamics of influenza viruses show strong genetic drift and purifying selection; positive selection has rarely been observed [13, 14, 53, 54]. A similar dichotomy is true for SARS-CoV-2 at within-host and population scales [55–58]. Our data suggest that these scales may be linked by a transmission process in which selective forces have an opportunity to act before potent stochastic forces within hosts. For example, antibodies present at mucosal surfaces could act on an antigenically diverse incoming viral population, mediating antigenic selection. Only positively selected variants would then be available to pass through the stochastic bottleneck associated with population expansion. Experiments performed in pre-immune hosts are needed to formally test this model.

In summary, our data reveal a two-stage transmission process in which transfer between hosts can include a large and diverse viral population but early events in the recipient host impose a stringent and stochastic bottleneck. Transmission may therefore represent an opportunity for selection to operate in the earliest phases of infection before any single genotype sweeps the population. This effect is expected to have a strong influence on viral evolution and is likely relevant across diverse viral families.

## MATERIALS & METHODS

### Ethical considerations

All the animal experiments were conducted in accordance with the Guide for the Care and Use of Laboratory Animals of the National Institutes of Health. The studies were conducted under animal biosafety level 2 containment and approved by the IACUC of Emory University (PROTO201700595) for the guinea pig (*Cavia porcellus*). The animals were humanely euthanized following guidelines approved by the American Veterinary Medical Association.

### Cells

Madin–Darby canine kidney (MDCK) cells were a gift from Dr. Daniel Perez, University of Georgia, Athens, GA. A seed stock of MDCK cells at passage 23 was amplified and maintained in Minimal Essential Medium (Gibco) supplemented with 10% fetal bovine serum (FBS; Atlanta Biologicals) and Normocin (Invivogen). 293T cells (ATCC, CRL-3216) were maintained in Dulbecco’s Minimal Essential Medium (Gibco) supplemented with 10% FBS and Normocin. All cells were cultured at 37 °C and 5% CO_2_ in a humidified incubator. The cell lines were not authenticated. All cell lines were tested monthly for *Mycoplasma* contamination while in use. The medium for the culture of influenza A virus in MDCK cells (virus medium) was prepared by supplementing Minimal Essential Medium with 4.3% bovine serum albumin (BSA; Sigma) and Normocin.

### Generation of Pan/99 NA-BC plasmid

The region of Pan/99 into which the barcode was inserted was first identified by aligning 593 sequences of H3N2-subtype influenza A viruses isolated between 1994 and 2004. A region with many nucleotide substitutions was identified in the NA segment from nucleotide position 484 to 532. Twelve synonymous mutations were identified within this region. Double-stranded Ultramers (IDT) were designed that contained degenerate bases with two possible nucleotides at each of the twelve chosen barcode sites (cacagtacatgataggaccccttaycgraccytattgatgaatgarttrggtgtyccattycayytrggracyaagcaagtgtgtatagca tggtcc).

Site-directed mutagenesis was used to insert an Xho1 restriction site into the wild-type reverse genetics plasmid, pDP Pan/99 NA, prior to barcode insertion. This was done to give a means of destroying the parental template following barcode insertion. To generate a linearized template for barcode insertion, the following steps were performed. Xho1 digestion of the plasmid stock was followed by phosphatase treatment using rSAP (NEB) to dephosphorylate the cut ends of the plasmid. The plasmid was then amplified by PCR using primers that extend outward from the barcode region: P99_NA_536F 5’-caagtgtgtatagcatggtcc-3’ and P99_NA_479R 5’-gggtcctatcatgtactgtg-3’. PCR purification (Qiagen QIAquick PCR Purification Kit) was used to isolate the linearized PCR product, followed by a dual digestion with Dpn1 and Xho1 to remove residual WT plasmid. PCR purification was repeated, and then an assembly reaction using the NEBuilder HiFi DNA Assembly Kit (NEB) was performed to insert the Ultramers into the linearized vector and re-circularize. The product was then transformed into DH5-α cells (NEB).

After plating onto LB-amp plates, approximately 1 x 10^4^ colonies were collected and pooled into LB-amp culture media and then incubated at 37°C for five hours prior to harvesting the bacterial population for plasmid purification by maxiprep (Qiagen Plasmid Maxi Kit). The presence of a diverse barcode in the plasmid stock was verified by next generation sequencing (see below). This plasmid stock was then used to generate Pan/99 NA-BC virus by reverse genetics in combination with seven plasmids encoding Pan/99 WT gene segments in a pDP2002 vector [59].

### Viruses

Pan/99 WT and Pan/99 NA-BC viruses were derived from influenza A/Panama/2007/99 (H3N2) virus (Pan/99 WT) and were generated by reverse genetics. In brief, 293T cells transfected with eight reverse genetics plasmids 16–24 h previously were co-cultured with MDCK cells in virus medium at 33°C for 40 h. Recovered virus was propagated in MDCK cells to generate working stocks. Propagation was carried out from low MOI to avoid accumulation of defective viral genomes but with a sufficient viral population size to maintain barcode diversity. Titration of stocks and experimental samples was carried out by plaque assay in MDCK cells. The presence of a diverse barcode in the virus stock was verified by next generation sequencing (see below).

### Growth kinetics

Replication of Pan/99 WT and Pan/99 NA-BC was determined in triplicate culture wells. MDCK cells in six-well dishes were inoculated at an MOI of 0.01 PFU/cell in PBS. After 1 h incubation at 33°C, inoculum was removed, cells were washed 3x with PBS, 2 mL virus medium was added to cells, and dishes were returned to 33°C. A 120 µL volume of culture medium was sampled at the indicated times points and stored at −80 °C. Viral titers were determined by plaque assay on MDCK cells.

### Guinea pig infections

Female Hartley strain guinea pigs weighing 250–350 grams were obtained from Charles River Laboratories and housed by Emory University Department of Animal Resources. Guinea pigs were sedated with ketamine (30 mg/kg) and xylazine (4 mg/kg) by intramuscular injection prior to intranasal inoculation or nasal lavage. Virus inoculum was given intranasally in a 300 μL volume of PBS containing 5 x 10^4^ PFU of Pan/99 NA-BC. At day 1 post-inoculation, one naïve animal was placed with each inoculated animal in either a cage that allowed for direct physical contact (to model transmission by all modes) or a cage in which the two animals were separated by a double-walled, perforated metal barrier (to model transmission via airborne infectious respiratory particles). Nasal lavage was performed with 1 mL PBS per animal on days 1-7 post-inoculation for inoculated animals and days 2-8 post-inoculation for exposed animals. Collected fluid was divided into aliquots and stored at -80 °C. Viral titers of nasal lavage samples were subsequently determined by plaque assay.

For the dose escalation experiment (**Supplemental Figure 5**), guinea pigs were obtained and sedated as described above. Virus inoculum was given intranasally in a 30 µL volume of PBS containing 1x10^2^ PFU, 3.3x10^2^ PFU, 1x10^3^ PFU, 3.3x10^3^ PFU, 1x10^4^ PFU, or 5 x 10^4^ PFU of Pan/99 NA-BC. Nasal lavage was performed with 1 mL PBS per animal on days 1-4 post-inoculation, and samples were stored as described above.

### Sample processing for next generation sequencing of barcode region

The following method was used to validate plasmid and virus stocks and to evaluate experimental samples. Viral RNA extraction (QiaAmp Viral RNA kit, Qiagen) was performed using a 140 µL volume of each nasal lavage sample or virus stock, followed by reverse transcription (Maxima RT, Thermo Fisher) with pooled Univ.F(A)+6 and Univ.F(G)+6 primers [41, 60]. Either cDNA or, for the purpose of plasmid validation, plasmid DNA was subjected to PCR amplification with primers flanking the barcode region. Samples were sent for amplicon sequencing to either Azenta Life Sciences or the Emory National Primate Research Center (ENPRC) Genomics Core. For samples sent to Azenta, PCR to amplify the region containing the barcode was performed using PFU Turbo AD (ThermoFisher) with primers with partial sequencing adapters that generated a 155-nt product (P99_NA_adptr_428-449_F 5’-acactctttccctacacgacgctcttccgatctggaacaacactaaacaacaggc-3’ and P99_NA_adptr_563-582_F 5’-gactggagttcagacgtgtgctcttccgatctgcttttccatcgtgacaact-3’). For samples sequenced by ENPRC, amplicon PCR was performed in a similar manner but using primers with sequencing adapters that generated a 100-nt product (P99NA_462F5_adptr 5’-tcgtcggcagcgtcagatgtgtataagagacagggactcctcagtacatgataggacccctt-3’ and P99NA_553Rev_adptr 5’-gtctcgtgggctcggagatgtgtataagagacagccatgctatacacacttgct-3’). Thirty PCR cycles were performed for all samples. Column-based PCR purification was performed on all samples followed by quantification of DNA using either NanoDrop or Qubit. Samples were normalized to 20 ng/uL in nuclease-free H_2_0.

Samples submitted to Azenta underwent Amplicon-EZ sequencing, an Illumina-based sequencing service compatible with amplicons of 150-500 nt in length that does not include a fragmentation step in library preparation. Amplicons were sequenced as 2 × 250 bp paired reads and demultiplexed prior to delivery. At ENPRC, library preparation was performed with the omission of a tagmentation step and sequencing was performed on an Illumin NovaSeq 6000 platform. Amplicons were sequenced as 2 x 100 bp paired reads and demultiplexed prior to delivery.

### Analysis of barcode sequencing data

Sequences were processed using our custom software, BarcodeID, available in the GitHub repository: https://github.com/Lowen-Lab/BarcodeID. Briefly, BarcodeID uses BBTools [35] to process raw reads, then uses a custom Python script to screen and identify barcode sequences present in each sample, then performs an error correction step to remove spurious barcode sequences, before finally calculating diversity statistics and writes summary tables. BBMerge screens for adapter sequences and merges forward and reverse reads using default settings, and BBDuk merges reads with a low average quality (<30). BarcodeID then screens each read to verify that the nucleotides at barcode and non-barcode sites match the nucleotides expected at those sites. Reads with mismatches are excluded from overall barcode sequence counts, but BarcodeID collects all high-quality variant amplicons and calculates overall mismatch rates by site to determine if any mutants with non-barcode alleles are driving any observed barcode dynamics. Samples with evidence of non-barcode driven dynamics were excluded from further analyses.

The error correction step was created to emulate the error correction software designed for bacterial 16S rRNA amplicons, DADA2 [61]. The largest difference between the two methods is how the nucleotide substitution model is generated. Where DADA2 uses a complicated statistical method, we can measure it directly from the backbone sites which should only deviate from the expected sequence due to mutation or any error accumulated through the process of library preparation and sequencing. Specifically, it measures the rate at which bases with a specific quality score match or do not match the expected base at each backbone site (λ_ji,Q_). For example, the rate of A to T with a quality score of 25 (λ_TA,25_) is the number of times a T with quality score of 25 was observed at a backbone site where A was expected, divided by the number of observations of all bases with a quality score of 25 at those sites. Using this error model and the observed prevalence of barcode sequences, BarcodeID estimates the probability that any barcode sequence could be derived from another through error. This probability calculated by:

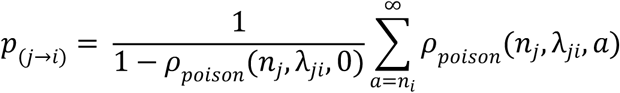

Where the probability of a barcode sequence *j* becoming barcode sequence *i* is Poisson distributed, and determined by the prevalence of both barcodes, *n_j_* and *n_i_*, and the product of probabilities of observing all nucleotide changes required to mutate barcode *j* into *i*.

BarcodeID uses the same agglomerative clustering process as DADA2 that attempts to iteratively identify barcodes that are too abundant to be explained due to error and collapse erroneous barcodes into larger barcode clusters. To initiate the agglomeration process, the most abundant barcode sequence with its associated quality score values in a sample is chosen as the initial seed barcode. BarcodeID then calculates the probability that every observed barcode sequence-quality combination could have been observed through mutation of the initial seed sequence (*p*_i,j_ where *j* is the seed barcode and *i* is the query barcode). Barcodes are then ranked in order of increasing probability of observation due to error. If the highest ranked barcode is higher than a Bonferroni adjusted p-value threshold, then it is used to create a new barcode cluster seed. This iterative process of adding the top seed before re-calculating the probability of observation due to error of all remaining barcodes relative to the growing list of seed barcodes continues until no more new seeds are found. Then, BarcodeID enters the agglomeration phase, where query barcodes are sorted in decreasing order of probability of observation due to error, and the barcode with the highest probability of being an error derived from a seed barcode is added to that seed barcode’s cluster. This process proceeds again iteratively until no barcodes can be collapsed into existing barcode clusters, at which point the remaining barcodes are included as new seeds until all observed barcodes are included. Importantly, the P_i>j_ of barcodes that are observed once will always be equal to 1 and are therefore unable to be clustered. Accordingly, singleton barcodes are excluded from calculating diversity statistics.

To account for uneven sampling intensity, BarcodeID performs repeated subsampling and linear regression to calculate diversity statistics that are more directly comparable between samples. Specifically, samples were subsampled at up to 19 different read depths (between 10^3 to 10^6 reads, where possible), with five iterations per subsampled depth. The median value of each diversity statistic at each depth was then used to fit the relationship between each diversity statistic and sampling intensity. The fit was then used to calculate an adjusted value for all samples at an equivalent depth. For alpha diversity statistics (richness, Shannon diversity, Simpson diversity, and evenness) the equivalent depth was 10,000 reads per sample because it provided the sufficient depth to distinguish samples across multiple levels of diversity, while minimizing bias due to variable sampling intensity. For beta diversity statistics (Bray-Curtis and Jaccard dissimilarity), 50,000 reads per sample was chosen because the added depth improved resolution in pairwise sample comparisons.

### Validation of barcode sequencing data

Samples were processed for barcode sequencing if infectious virus could be detected therein by plaque assay (which has a limit of detection of 50 PFU/mL). We found that many reads were obtained and analyzed for plaque-positive samples, irrespective of viral titer. To evaluate the reproducibility of sequencing results, samples from three guinea pigs were sequenced in triplicate. This set of samples includes high titer, low titer and plaque negative samples. For each sample, the original RNA extract was processed independently three times to produce, amplify, purify and sequence the cDNA. Results indicate that barcode composition detected in plaque-positive samples is highly reproducible, while barcode composition detected in the plaque-negative samples is not reproducible (**Supplemental Figure 4C**).

### Preparation of nasal lavage samples for viral whole-genome sequencing

Viral RNA extraction (QiaAmp Viral RNA kit, Qiagen) was performed using a 140 µL volume of each nasal lavage sample, followed by one-step reverse transcription PCR amplification of full viral genomes using pooled Univ.F(A)+6, Univ.F(G)+6 primers and Univ.R primers and SuperScript III Platinum kit (ThermoFisher) [41, 60]. Following PCR purification (Qiagen QiaQuick PCR Purification Kit), cDNA was processed at the ENPRC core for sequencing on an Illumina NovaSeq 6000 platform. Samples were sequenced as 2 x 100 bp paired reads and demultiplexed prior to delivery.

### Analysis of whole-genome sequencing data

Whole genome sequencing reads were merged and filtered for low average quality (≥30) using BBMerge, then separated according to the segment using BLAT [62]. The reads were then mapped to their corresponding reference segment using BBMap, with local alignment set to false. From these alignments, we used custom Python scripts to identify iSNVs and reads that map to the barcoded region of the NA segment. Cutoffs for inclusion of iSNVs were set empirically. First, sites were evaluated based on their total coverage and the average quality and mapping statistics. Only sites with ≥100x coverage were considered. For minor variants at these sites to be included in subsequent analyses, they were required to be present at ≥1% frequency and have an average phred score of ≥35, and the reads that contained the minor allele at any given site also had to have sufficient mapping quality to justify inclusion. Specifically, reads containing the minor allele needed an average mapping quality score of ≥40, the average location of the minor allele needed to be ≥20 bases from the nearest end of a read, and the reads overall needed to have ≤2.0 average mismatch and indel counts relative to the reference sequence.

### Analysis of beta diversity

Dissimilarity between two populations can be measured using beta diversity metrics, and in this study, we used the both Jaccard index and the Bray-Curtis dissimilarity [63, 64]. Both metrics consider the species shared between two populations, and in this case, unique barcodes were considered species. In addition to presence/absence data, the Bray-Curtis dissimilarity also reflects abundance data. For either measure, a value closer to one indicates that the two populations are more dissimilar, whereas a value closer to zero indicates that the two populations are more alike in composition. Pairwise comparisons of barcode data were made between the viral populations present in plaque-positive nasal lavage samples acquired from a contact pair of guinea pigs.

## Data availability

All data generated or analyzed during this study are included in the manuscript. At the time of initial submission, all raw sequencing data has been uploaded to NCBI’s Sequence Read Archive under the accession number PRJNA1253752.

## Acknowledgments/Funding

We are grateful to Katia Koelle for helpful discussion. This study was supported in part by R01 AI165644 and the National Institute of Allergy and Infectious Diseases (NIAID) Centers of Excellence for Influenza Research and Response (CEIRR) contract no. 75N93021C00017.

---

**Supplemental Figure 1.**
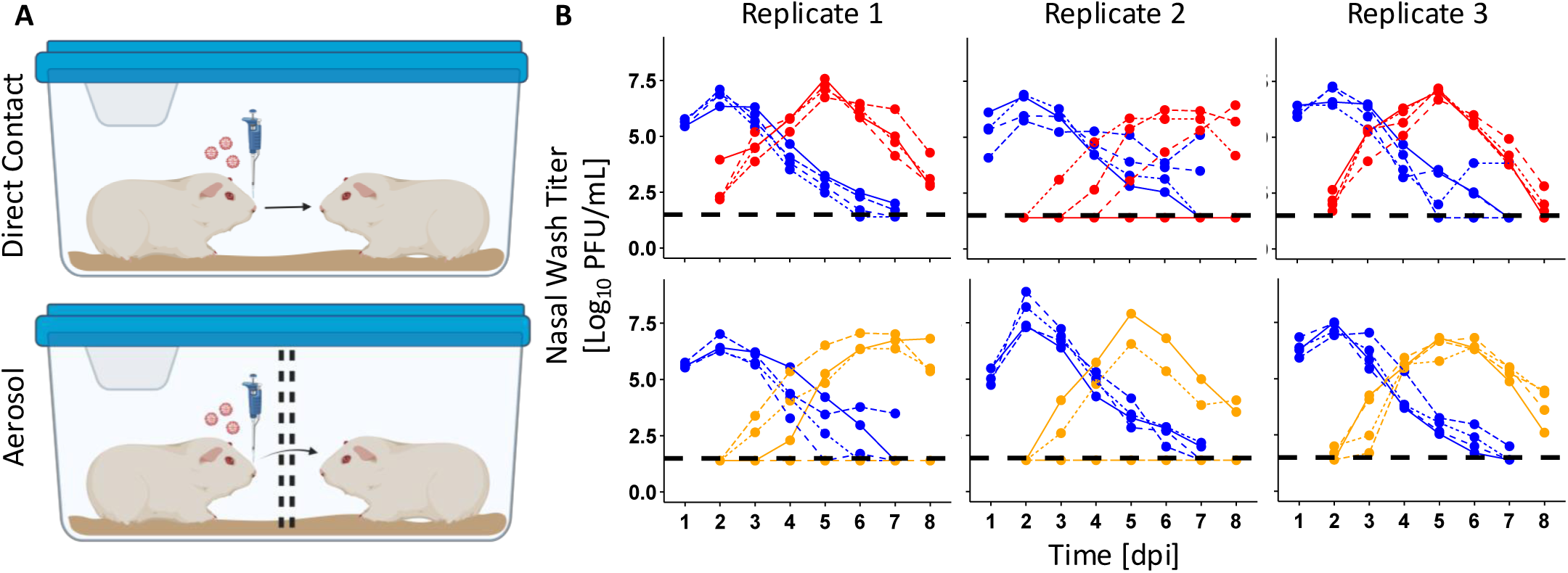
Pan/99 NA-BC infects inoculated animals and transmits to exposed animals. **A)** Schematic showing transmission set-up in guinea pigs. For each of the three experimental replicates, eight guinea pigs were intranasally inoculated with 5 x 10^4^ PFU of Pan/99 NA-BC in 300 μL. Twenty-four hours post-inoculation, a single naïve animal was placed with each inoculated animal. Cages either allowed for direct contact (n = 4) or maintained separation with a double-walled, perforated metal barrier (n = 4). **B)** Viral titers in nasal lavage samples in direct contact and aerosol exposure settings from replicates 1, 2, and 3. Inoculated animals are shown in blue and exposed animals in red (direct contact) or yellow (aerosol contact). The dashed black line represents the limit of detection (50 PFU/mL). Paired animals share the same line type. Supplemental Figure 1A created in BioRender.

**Supplemental Figure 2.**
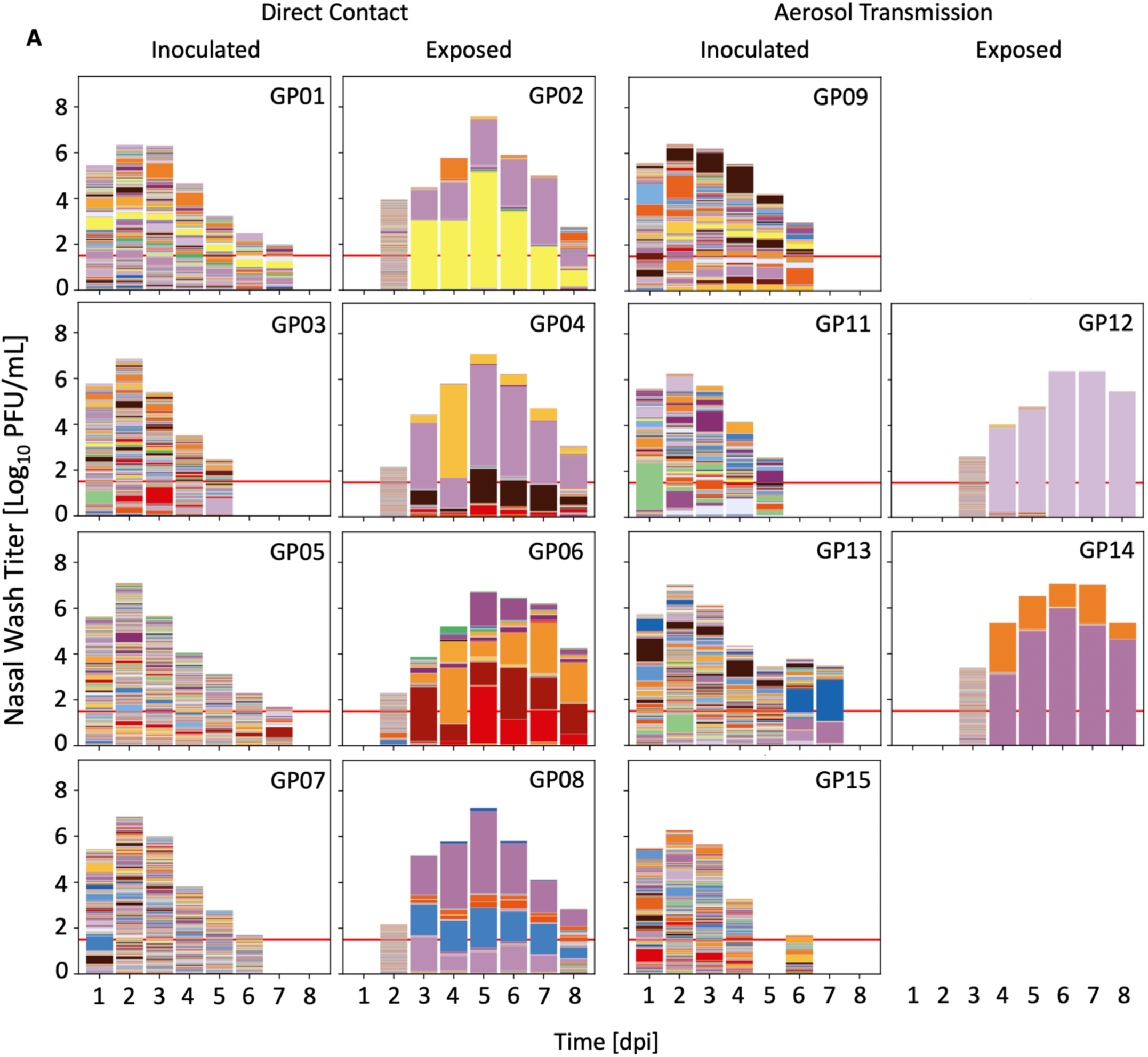
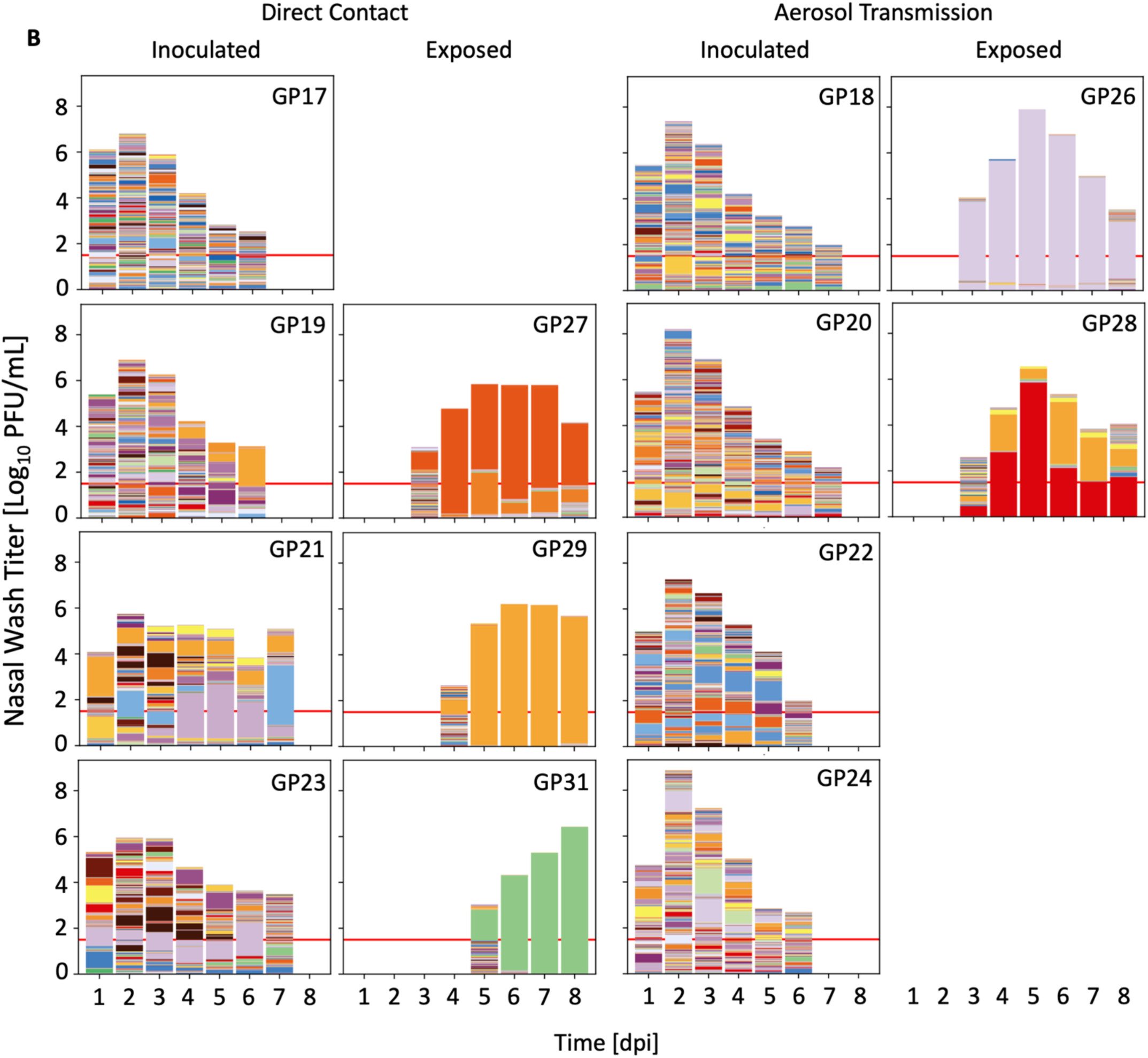
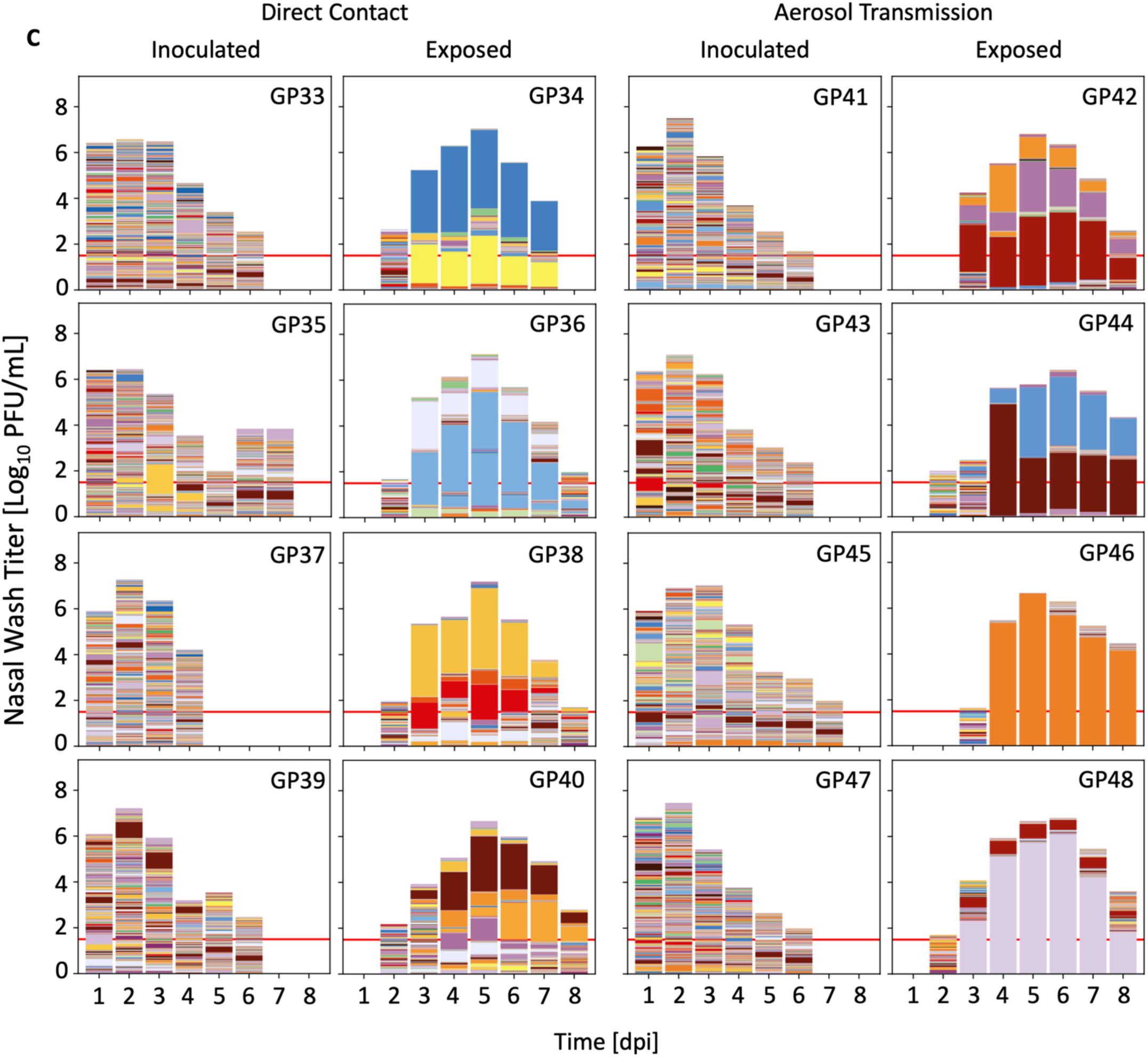
Population diversity declines between inoculated and exposed guinea pigs. Data from experimental replicates 1, 2, and 3 are shown in panels **A, B,** and **C,** respectively. Guinea pig ID numbers are shown in the upper right corner of each plot. Nasal lavage titers are indicated by the total height of each bar. Colors within the bars represent unique barcodes, and the height of each color indicates the relative frequency within the sample. Only samples that were plaque-positive are shown. Red lines show LOD of 50 PFU/mL. Plots for individual animals are paired with those of their cage mate. For the exposed animal in the first aerosol transmission pair of Replicate 1 (GP10), most reads were discarded due to poor quality, and the data from this animal were excluded from further analyses.

**Supplemental Figure 3.**
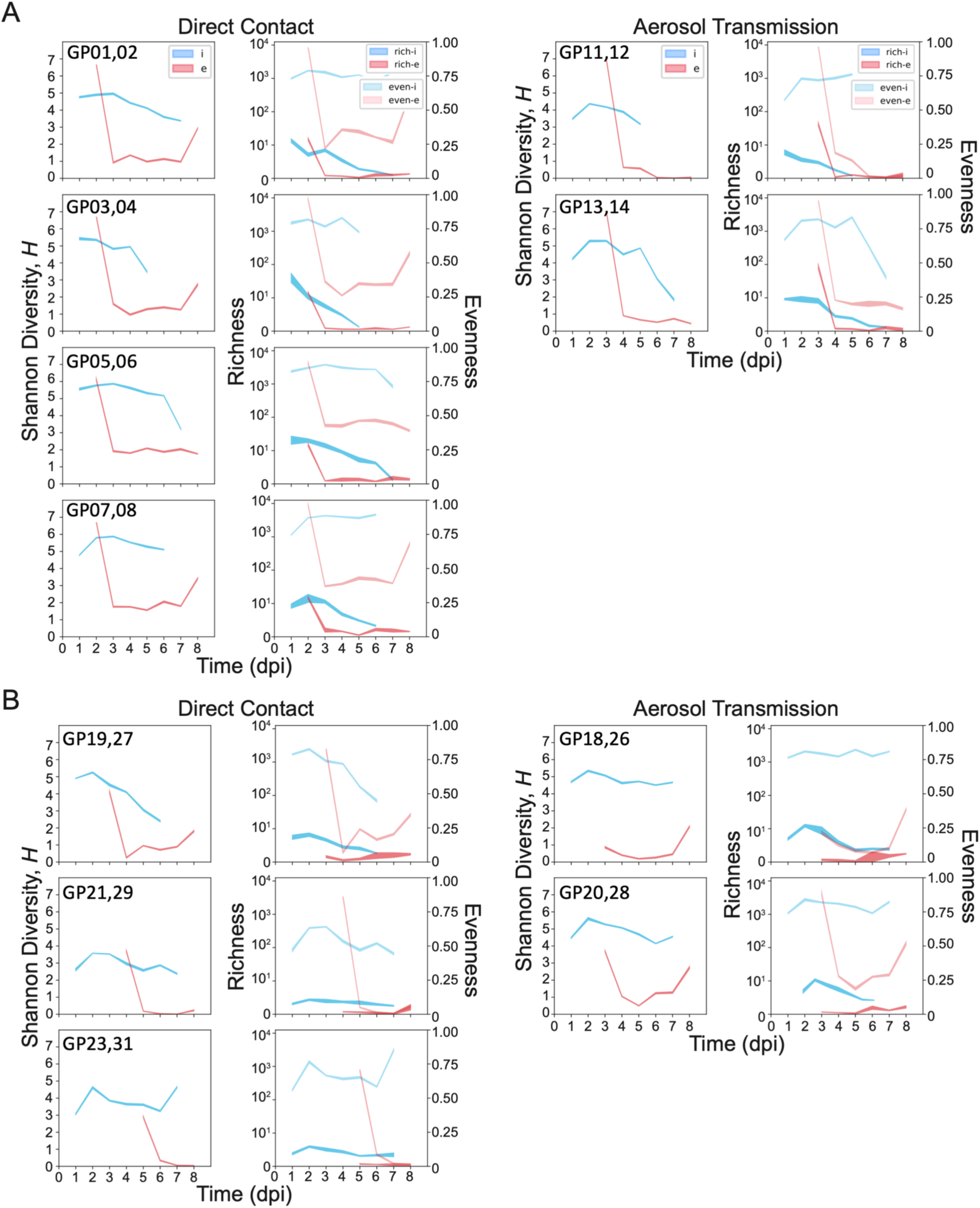
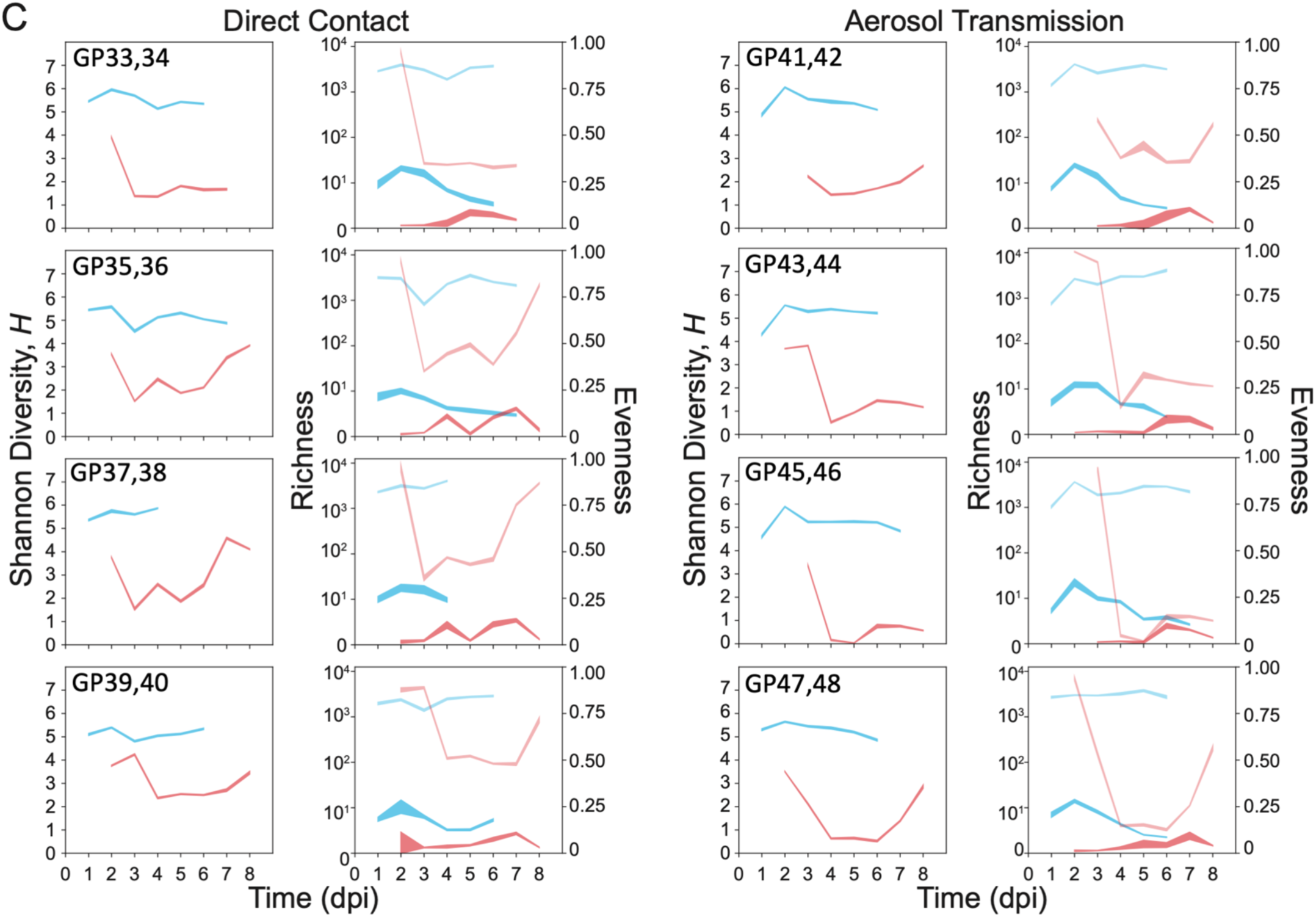
Changes in richness and evenness both contribute to alterations in diversity. Shannon diversity (left), richness (right, left axis), and evenness (right, right axis) were determined for inoculated animals (i, blue) and exposed animals (e, red) in replicate experiments 1, 2, and 3 (**A, B,** and **C**, respectively). Data from cage mates are paired. In the right-hand plots, bold colors show richness and faded colors show evenness. Line widths show 95% confidence intervals based upon subsampling of barcode reads.

**Supplemental Figure 4.**
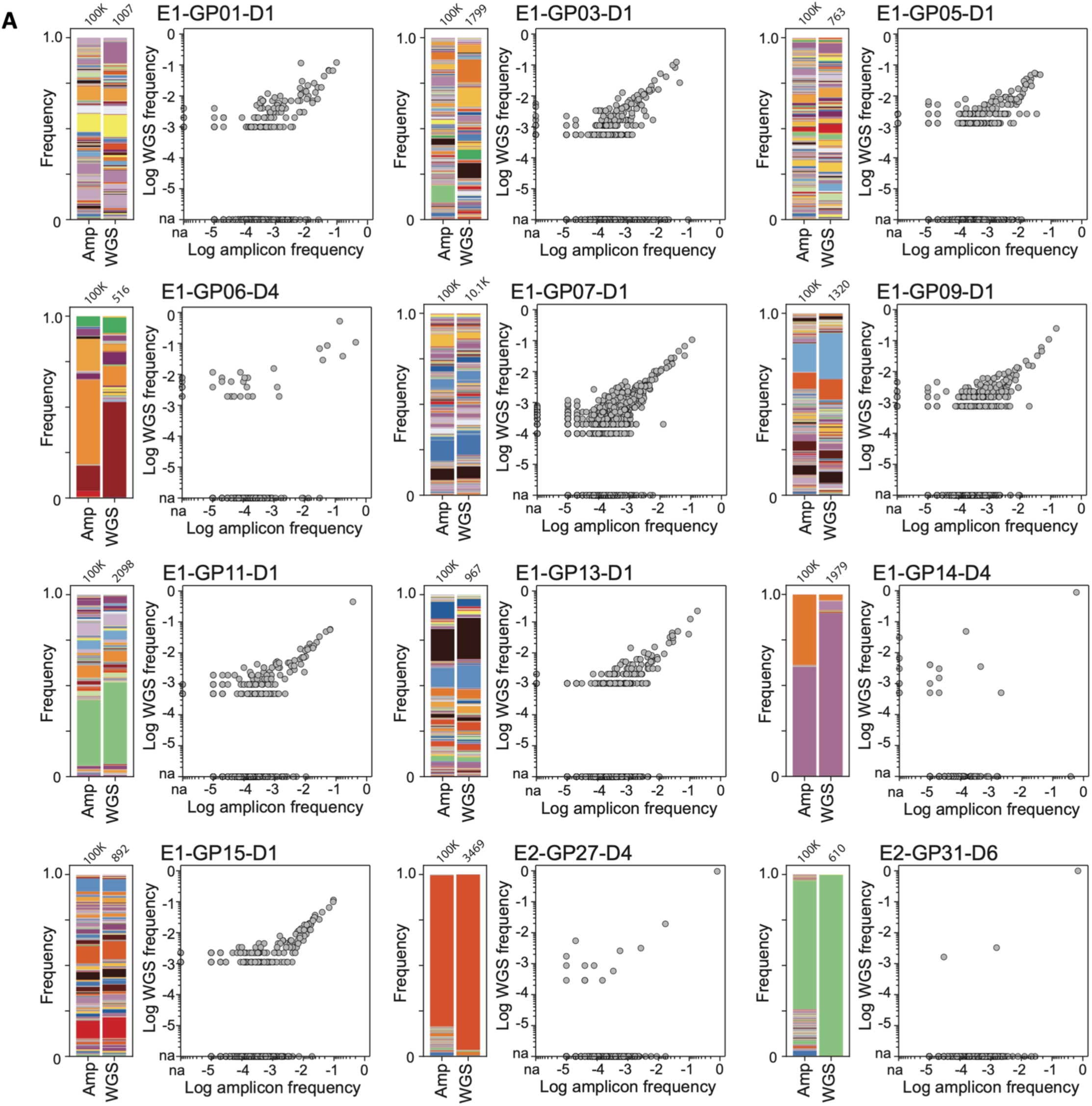
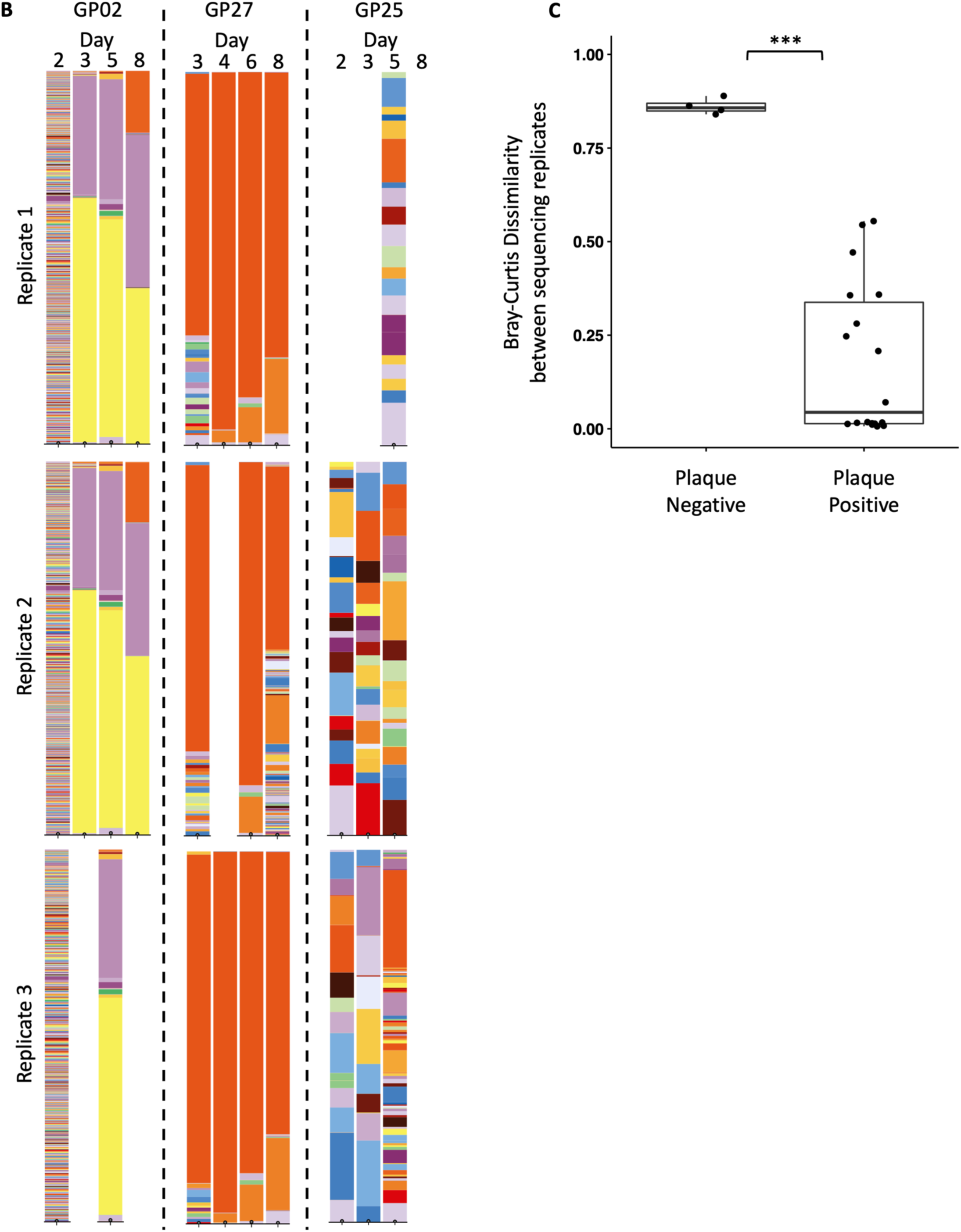
Targeted amplicon sequencing of the barcode region is reliable and reproducible. **A)** Whole-genome sequencing (WGS) was performed on a subset of samples from experimental replicates 1 and 2. WGS reads that spanned the length of the barcode region were compared to targeted barcode sequencing (amplicon or Amp) results for each sample. Each graph is labeled with the experiment number (E1, E2, or E2), the guinea pig number (GP01, GP03, etc.), and the day post-inoculation (D1, D4, etc.). The number of reads mapping to the barcode region is indicated above the stacked bar plots. **B)** A subset of positive nasal wash samples from two animals (GP02, GP27) were sequenced in triplicate to evaluate reproducibility of sequencing results. These were compared to an animal that was plaque-negative throughout the experiment (GP25). **C)** Bray-Curtis Dissimilarity values between replicate sequencing runs from each sample are plotted for the plaque-negative animal and the plaque-positive animals (*** p-value < 0.0005, Wilcoxon Sign-Rank test).

**Supplemental Figure 5.**
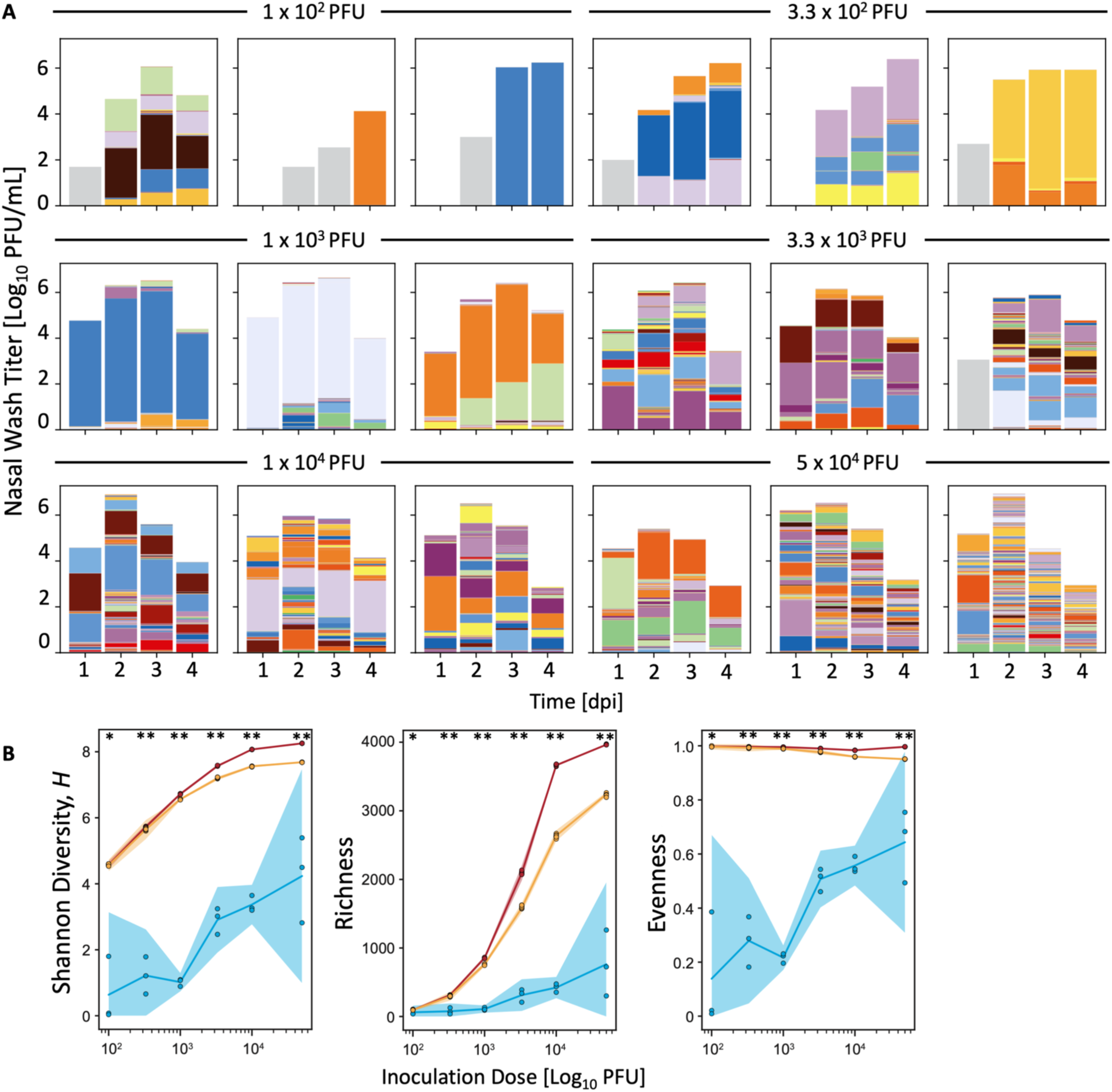
Growth-induced bottlenecks are not detected in inoculated animals, even at low inoculation doses. At each inoculation dose indicated above the plots, three guinea pigs were inoculated intranasally with Pan/99 NA-BC. **A)** Nasal lavage titers and barcode compositions over time. The height of the bars indicates viral titer in samples above the limit of detection (50 PFU/mL). Colors indicate barcodes detected, with barcode frequency shown by the height of the color. **B)** Maximum measured Shannon diversity, richness, and evenness of barcode compositions in each animal are plotted by inoculation dose (blue). For comparison, theoretical data are plotted to show the expected characteristics of inocula of each size derived from a perfectly even population of 4096 barcodes (red) and from the Pan/99 NA-BC passage 1 stock (orange). 95% confidence intervals are shown with shading. Statistical significance was determined at each inoculation dose by Kruskal-Wallis Test (* p-value < 0.01, ** p-value < 0.001) comparing the medians of the experimental data with those of the perfectly even population and that of the passage 1 viral stock.

